# Subacute and Chronic Cognitive and Cerebrovascular Functional Consequences of Mild Traumatic Brain Injury in Rats

**DOI:** 10.64898/2026.07.22.740068

**Authors:** Jonathan Lifshitz, Ashley Ruhland, Jackson Bisesi, Nina Karamanova, L. Matthew Law, Daniel R. Griffiths, Alberto Fuentes, Maurizio Bergamino, Connor Leighty, Thomas L. Broderick, Camelia Burciu, Taben M. Hale, Ashley Stokes, Raymond Q. Migrino

**Affiliations:** Phoenix Veterans Affairs Health Care System, Phoenix Arizona, USA; University of Michigan, Ann Arbor Michigan, USA; Ann Arbor Veterans Affairs, Ann Arbor Michigan, USA; Barrow Neurological Institute, Phoenix Arizona, USA; Midwestern University, Glendale Arizona, USA; Alabama College of Osteopathic Medicine, Dothan Alabama, USA; University of Arizona College of Medicine-Phoenix, Phoenix Arizona, USA

## Abstract

Traumatic brain injury (TBI) is the main cause of death and disability in the United States in people younger than 35 years old and is a common cause of wartime injuries. Mild TBI (mTBI) is a predisposing factor for later development of dementia and cerebrovascular disease. Aerobic exercise was reported to improve cognitive function in chronic TBI through improved vascular function. The aims of the study are to characterize and correlate the subacute (10 weeks) and chronic (12 months) cognitive and cerebrovascular functional impairment following mTBI and evaluate the modulating effect of exercise early or late following mTBI on these changes. Sprague-Dawley rats received midline fluid percussion injury or sham procedure and followed for 10 weeks or 12 months with a subgroup of mTBI rats undergoing 5-week treadmill aerobic exercise 2 weeks (early) or 10 months (late) post-injury for 6 weeks. Cognitive function was assessed using novel object recognition (NOR) and novel object location (NOL) tests. Regional cerebral blood volume (CBV) and cerebrovascular reactivity following hypercapneic stimulation (CVR) using contrast magnetic resonance imaging (MRI) and ex vivo pial artery vasoreactivity to intraluminal pressure, angiotensin II and diethylenetriamine NONOate (DETA NONOate) were measured. There was no difference in NOR or NOL at 10 weeks between mTBI and sham. NOR, but not NOL, was reduced in mTBI rats at 12 months. CBV at 10 weeks was higher in the primary somatosensory trunk cortex (trunk) and dentate gyrus regions in mTBI, but not at 12 months. CVR was lower in the trunk region of mTBI rats at 12 months. Compared to sham response at 10 weeks, there was impaired arterial constriction response to 90 mm Hg intraluminal pressure in mTBI rats at 10 weeks and in sham and mTBI rats at 12 months, with no difference seen in response to angiotensin II or DETA-NONOate exposure. There was no correlation between cognitive and vascular outcomes at 10 weeks or 12 months. Early or late exercise did not affect 12-month cognitive or vascular function following mTBI. The study showed chronic impairment in short-term memory cognitive function, regional cerebrovascular reactivity and early onset of impaired cerebrovascular myogenic response in rats subjected to mTBI. The findings of persistent cognitive and cerebrovascular impairment in this animal model enhance our understanding of the long-term consequence of mTBI.

## Introduction

Approximately 3 million patients with traumatic brain injury (TBI) seek medical attention in the United States annually^1,2^. TBI is reported in 8-22% of military personnel involved in combat operations^2,3^ with explosion or blast injury being the most common cause of wartime injuries during the war in Iraq and Afghanistan^4^. TBI is also the main cause of death and disability in people younger than 35 years old in the United States^5^. Traumatic brain injury (TBI) occurs when the skull is exposed to an impact or mechanical force which results in both a primary and secondary insult to the brain ^6–8^. The primary insult is characterized by an initial impact to the skull and the brain, while the secondary insult is characterized by neuronal and vascular damage, apoptosis, inflammatory processes, and proteolytic pathways. Secondary damage continues to develop even after the initial impact.

Chronic neurologic and cardiovascular adverse outcomes are associated with TBI. Cognitive dysfunction is a long-term sequela of TBI with deficits in attention, memory and executive functioning^5,9^. Epidemiologic data indicate that individuals with TBI history have a higher risk of developing dementia^10,11^. Among World War II veterans, nonpenetrating head injury in early adulthood was associated with increased risk of Alzheimer’s disease and other dementias (AD/ADRD)^12^. Additionally, registry-based studies demonstrated increased risk of cardiovascular and cerebrovascular disorders in the chronic phase of TBI recovery^13^. Both mild and moderate-severe TBI were associated with increased risk of ischemic stroke or transient ischemic attack within a median onset of 3.49 years after TBI^14^. TBI and dementia disorders such as AD/ADRD share a common pathology of vascular dysfunction. Patients ∼5 years following mTBI injury were reported to have hypoperfused regions in the frontal, prefrontal, temporal cortices as well as subcortical structures with regional hypoperfusion concordant with neuropsychological localization^15^. Retired boxers showed cerebral hypoperfusion coupled with neurocognitive dysfunction^16^. Epidemiologic, preclinical and clinical data similarly show that vascular disease is strongly associated with dementia and that vascular dysfunction leading to cerebral hypoperfusion is critical in the early stages of AD^17^.

While moderate and severe TBI acutely lead to significant irreversible or partly reversible neuronal and vascular damage^18,19^ that persist to chronic neurovascular deficits, the mechanisms by which mild TBI (mTBI), which represents the great majority (60-95%) of TBI cases^20,21^, cause long-term cognitive and cerebrovascular dysfunction remain poorly understood. This knowledge gap is exacerbated by the lack of preclinical animal model studies with sufficiently long follow-up of chronic neurovascular consequences of mTBI. We previously showed that rats subjected to mTBI using midline fluid percussion injury (FPI) had impaired cognitive function and altered smooth muscle-dependent cerebral vasoreactivity at 6 months post-injury^22^. To further understand the long-term neurovascular pathology of mTBI and its underlying physiologic mechanisms, we compared cognitive function, *in vivo* cerebrovascular function and *ex vivo* cerebrovascular vasoreactivity at 10 weeks and 12 months in rats subjected to FPI or sham injury. Aerobic exercise has been shown to improve cognitive function in various disease models such as stroke and neurodegenerative conditions through improved cerebral autoregulation, neuroplasticity and neurogenesis, but there have been few studies in TBI^23^. Additionally, we sought to evaluate whether aerobic exercise early or late following mTBI would reduce mTBI-induced cognitive and vascular dysfunction.

## Methods

### Animals

The study was approved and supervised by the Phoenix Veterans Affairs/Barrow Neurological Institute at St. Joseph’s Hospital and Medical Center Institutional Animal Care and Use Committee (IACUC), which served as the IACUC of record. The University of Arizona served as the operational IACUC. Male (N =44) and female (n = 44) Sprague-Dawley rats (8-10 weeks old, 184-440g, Charles River Laboratories, Wilmington, MA) were acclimated for one week in their home cages. Rats were housed in a reverse-light cycle environment with access to standard rat chow and water *ad libitum*. Adequate measures were taken to limit pain and discomfort. Experiments were conducted in accordance with the University of Arizona, RIGOR and Animal Research: Reporting of *In Vivo* Experiments (ARRIVE) guidelines for the use of laboratory animals. In the conduct of these experiments, 26 rats (11 male; 15 female) died as a result of surgery or injury before group assignment.

### Midline fluid percussion injury

After 1 week of acclimation, rats underwent surgery for midline fluid percussion injury (FPI; 1.85-2.05 atm) or the uninjured sham. All animals were randomly separated into cohorts of uninjured (sham) and brain-injured (TBI) animals. The researcher who performed the surgery was different from the one assessing cognitive function, *in vivo* imaging data, or *ex vivo* vascular function. Surgery was consistent with previously described methods ^22^. Animals were anesthetized with 5% inhaled isoflurane anesthesia and maintained at 2% isoflurane on a ventilator. Body temperature was maintained with a Deltaphase® isothermal heating pad (Braintree Scientific, Inc., Braintree, MA, USA). In a head-holder assembly (Kopf Instrument, Tujunga, CA, USA), a midline scalp incision exposed the skull. A 4.8mm trephine-derived craniotomy was then centered between bregma and lambda at the sagittal suture. The meninges and superior sagittal sinus were undisturbed throughout this process. An injury hub was created with removal of the needle from a Luer-Lok needle hub which was cut, beveled, and scored to fit within the craniotomy. A skull screw was secured in a 1-mm hand-drilled hole in the right frontal bone. The injury hub was affixed over the craniotomy with cyanoacrylate gel and methyl-methacrylate (Hygenic Corp., Akron, OH, USA) applied around the injury hub and screw. The incision was sutured at the anterior and posterior edges and topical bacitracin, and lidocaine ointment were applied. Animals were set in a clean cage placed upon a heated pad and monitored until ambulatory.

For injury induction, animals were re-anesthetized with 5% inhaled isoflurane approximately 60-90 minutes after conclusion of the surgical procedure. The dura was inspected through the injury-hub assembly, which was then filled with saline prior to fluid percussion device attachment (Custom Design and Fabrication, Virginia Commonwealth University, Richmond, VA). Animals were placed under anesthesia prior to injury though anesthesia was not maintained throughout injury induction. Animals received brain-injury by releasing the pendulum onto the fluid-filled cylinder. Upon induction, animals were monitored for the presence of a forelimb fencing response and the return of the righting reflex as indicators of injury severity ^24^. The injury hub was removed *en bloc*, dura integrity observed, bleeding controlled with saline and gauze, and incision site stapled. Brain-injured animals had righting reflex recovery times of 473.6 ± 133.5 sec (range 279 to 870 sec), with apnea ranging between 5 and 180 seconds. Twenty-six of 88 rats did not survive 24 hours after the surgery (11 male; 15 female). After recovery of the righting reflex, animals were placed in a warmed holding cage before being returned to the housing room. Each rat was evaluated for post-operative health for three days after surgery. Appropriate interventions, including but not limited to injection of subcutaneous bolus of saline and provision of wet food mash were made as clinically indicated. A group of rats constituted the subacute group and were sacrificed 10 weeks following surgery and a separate group of rats constituted the chronic group and were sacrificed 12 months after surgery.

### Aerobic Exercise

A subset of rats in the chronic mTBI group underwent aerobic exercise training starting at 2 weeks post-mTBI (early exercise) or 10 months post-mTBI (late exercise) for six weeks, including acclimation period. Exercise training consisted of treadmill running (Exer 3/6, Columbus Instruments, Columbus OH, USA) one session per day in the morning under red light conditions, 5 days a week. Prior to onset of training, rats were initially acclimated to daily 10-minute sessions of treadmill running for a period of one week. After this acclimation period, exercise training consisted of a graded incremental increase in duration and intensity starting at 15 meters/min for 15 min for week 1 and increasing to 17 meters/min for 20 min for week 2. For the remaining 3 weeks, rats ran at 17-19 meters/min for 25-30 minutes, corresponding to an estimated ∼46-50% maximal oxygen carrying capacity^25^.

### Cognitive function evaluation

Cognitive function assessment took place 10 weeks, 6 months and 12 months after surgery. Cognitive tasks took place in a square arena measuring 68.58cm x 68.58cm, while white noise played at ∼48dB to mask environmental noises. Ethovision software recorded animal position and movement during test sessions (Noldus, Leesburg, VA, USA). Behavioral Observation Research Interactive Software (BORIS, https://www.boris.unito.it/) allowed for event quantification. The novel object recognition (NOR) task assessed short-term memory ^26^. Rats acclimated to the arena for 2 minutes. Animals were then presented with “twin” objects (A1, A2) in opposite corners of the apparatus (5 minutes) in the training trial. After a 4-hour delay, rats were returned to the arena with one object (A1) having been replaced with a novel object (B1). Animals without short-term memory impairment were expected to explore the novel object (B1) more than the remaining familiar object^27^.The NOR test served as the novel object location (NOL) training trial, with a 24-hour latency between. The NOL task tested long-term spatial memory^28^. The novel object from the NOR test did not move location (B1), while the familiar object (A1/A2) was moved to the opposite corner of the arena. Healthy rats were expected to prefer the object in the novel location (A1/A2). The temporal order recognition (TOR) task assessed working reference memory through the ability to recognize the order of objects presented over time^28^. Animals were exposed to two training trials and a test trial. Five minutes were allotted to explore a pair of objects in training trial 1 (objects C1, C2) and training trial 2 (objects D1, D2), followed by a two-minute intertrial interval (ITI). Then one object from each trial (one object from set C1/C2 and one object from set D1/D2) was presented in the test trial. Training trials had ITIs of 3 minutes, with a 5 minute ITI before the test trial. Animals with intact reference memory should spend more time with the initial object (C1/C2) versus the recent object (D1/D2).

For all tests of cognitive function, active exploration time was determined as time spent within ∼2cm of the object. Animals were assessed for active exploration of both objects, and a discrimination index was determined with a formula of (time spent with object of interest/time spent with both objects). A discrimination index of 0.5 is equivalent to chance performance and comparable exploration of both objects^28^; discrimination index above 0.5 indicates expected cognitive performance.

### Magnetic resonance imaging

#### MRI acquisition

MRI data were acquired at 10 weeks (subacute group) or 12 months (chronic group) post-surgery using 7T (Bruker Biospec 70/30, Billerica, MA) using a 40-mm transmit/receive rat head volume coil (Bruker T13161V3, Billerica, MA). Anesthesia was induced using 3% isoflurane in air and maintained with 1–2.5% isoflurane in air. Respiration was monitored continuously, and body temperature was maintained at 38°C using warm circulating water. Pre-contrast anatomical images were acquired using a rapid acquisition with relaxation enhancement (RARE) sequence, with the following parameters: repetition time (TR) = 2505 ms, echo time (TE) = 46.7 ms, RARE factor = 8, field of view (FOV) = 40 × 40 mm², resolution = 0.267 × 0.267 mm², 20 slices, slice thickness = 1 mm, 4 averages. Multi-gradient-echo images for quantitative T2* mapping were acquired using the following parameters: TR = 1000 ms; 6 echoes; TE range = 4-24 ms; 4 ms echo spacing; flip angle = 45°; FOV = 35 × 35 mm²; resolution = 0.233 × 0.233 mm²; 6 slices; slice thickness = 1 mm; 6 averages. Following baseline imaging, iron-oxide contrast agent (monocrystalline iron oxide nanoparticle; MION) was administered via tail-vein catheter at a 10 mg/kg dose. Post-contrast multi-gradient-echo T2*-weighted images were acquired to quantify cerebral blood volume (CBV), using the same protocol described above. Animals were then exposed to carbogen gas (95% O₂ / 5% CO₂) for a 10-minute equilibration period; subsequently, post-contrast T2*-weighted images were acquired using the same imaging protocol under continuous carbogen administration. These images were used to assess cerebrovascular reactivity (CVR) under hypercapnic stimulation.

### MRI Image analysis

Image processing was performed using in-house software developed in MATLAB (MathWorks, Natick, MA, USA). Quantitative T2* maps were generated by fitting the multi-echo signal to a monoexponential signal decay model using nonlinear least-squares estimation. R2* maps were calculated as the reciprocal of T2* (R2* = 1/T2*). CBV (in ml/100 g tissue) was calculated from the change in R2* relaxation rates between the pre-contrast and post-MION acquisitions, as previously described^29^. CVR was calculated from the carbogen-induced change in R2* relaxation rates, normalized by the baseline CBV^30,31^. Region-of-interest (ROI) analyses were performed using the SIGMA rat brain anatomical atlas^32^. More specifically, the anatomical RARE images were resampled to an isotropic resolution of 0.15 mm^33^ and nonlinearly registered to the SIGMA standard template using the ANTs SyN algorithm^34^. The resulting inverse transformation matrix and deformation field were then applied to map the atlas ROIs into each subject’s native RARE space. The T2*-weighted images corresponding to the first TE were linearly registered to the corresponding RARE images, and the resulting transformation matrix was applied to the CBV and VR maps. All ROI measurements were extracted in native RARE space. The following ROIs were analyzed: cornu ammonis area 1 and 3 of the hippocampus (CA1 and CA3), dentate gyrus (DG), corpus callosum (CC), thalamus, striatum, primary somatosensory barrel field cortex (BF1), and primary somatosensory trunk cortex (Trunk) (Figure 1).

**Figure 1.**
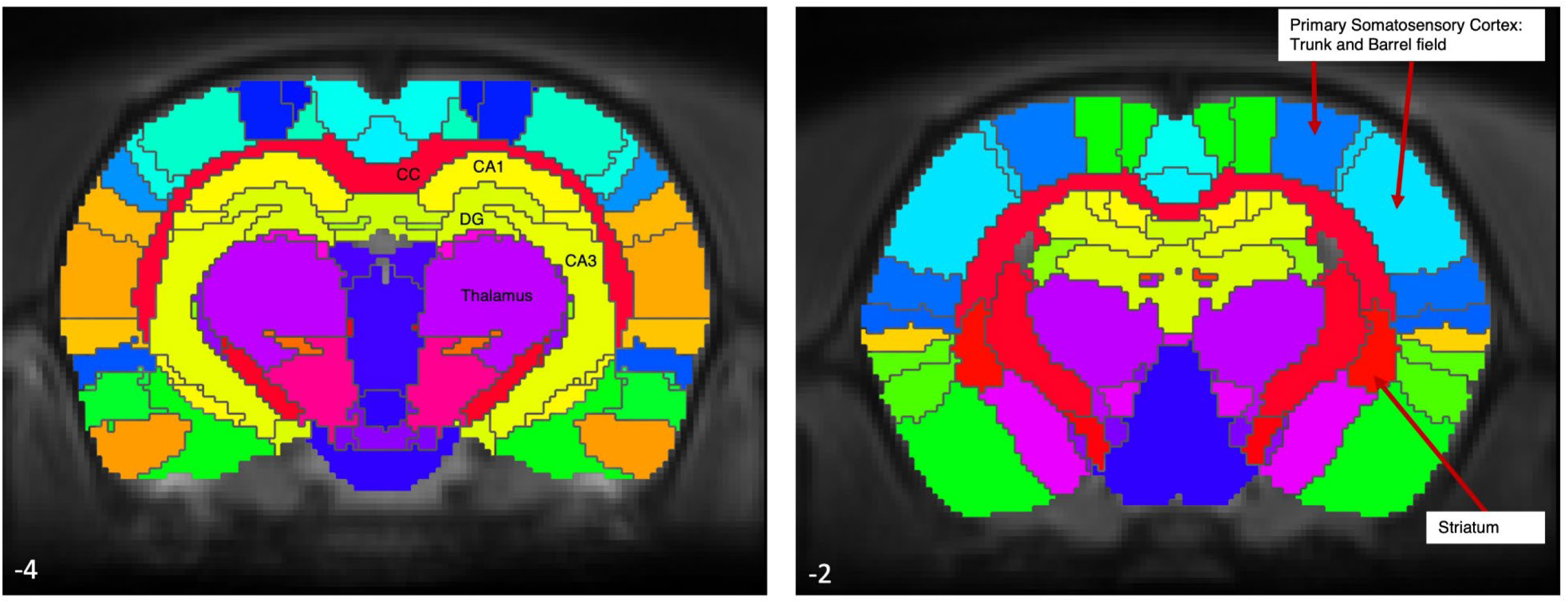
Atlas of brain regions analyzed with MRI. Cerebral blood volume and cerebrovascular reactivity were measured in the following select regions of interest with corresponding bregma levels: cornu ammonis area 1 and 3 of the hippocampus (CA1 and CA3), dentate gyrus (DG), corpus callosum (CC), thalamus, striatum, primary somatosensory barrel field cortex and primary somatosensory trunk cortex (trunk).

### Ex vivo cerebrovascular function assessment

Rats were euthanized either at 10 weeks (subacute group) or 12 months (chronic group) post-surgery via Euthasol (sodium pentobarbital and phenytoin, 340 mg/kg, Virbac Corporation, Westlake TX) overdose and circle of Willis (pial) arteries were carefully dissected from brain tissue and immediately placed in 4-(2-hydoxyethyl)-1-piperazineethanesulfonic acid (HEPES) buffer. The methods were previously described^22^ but with some modifications. Middle or posterior cerebral arteries were isolated and cannulated and arterial luminal diameters measured using videomicroscopes throughout the procedures. Myogenic tone response to intraluminal pressure was assessed by sequential intraluminal pressure exposure from 0 to 30, 60 and then 90 mm Hg (30 minute stabilization per increment). Myogenic response was reported as percentage change from luminal diameter at 0 mm Hg. Following stabilization at 90 mm Hg intraluminal pressure, the vessel was exposed to increasing doses of angiotensin II (10^-10^-10^-7^M, Sigma Aldrich, St. Louis MO, 4 min duration between doses), a potent cerebral arterial smooth-muscle dependent vasoconstrictor^35^. This was followed by diethylenetriamine NONOate (DETA NONOate, Cayman Chemicals, Ann Arbor MI) 10^-4^M, a nitric oxide donor to evaluate smooth muscle-dependent dilation^22^. Vasoconstrictor response to angiotensin II and DETA NONOate was reported as percentage change of luminal diameter from baseline pre-treatment diameter.

## Data and statistical analyses

Sample size considerations: Informed by cognitive function data from prior work^22^ and assuming the difference in NOR observed will be similar at 12 months (difference 0.26, standard deviation of 0.19 and 0.15), a sample size of at least 8 per group (sham or mTBI) were planned to allow us to show significant difference with α=0.05 and β=0.80. Pre-planned analysis was to combine data from male and female rats.

Data are expressed as means±standard error of means. Pairwise analyses were performed using Student’s t-test after confirming normal distribution of data. Group analyses were performed using one-way analysis of variance with post-hoc pairwise test using Tukey-Kramer and accounting for multiple comparisons. Correlation analyses were performed using Pearson correlation. Analyses were performed using MedCalc version 19.8 (MedCalc Software, Ostend, Belgium) and GraphPad Prism 10 (GraphPad Software, Boston MA). Significant p value was set at p<0.05 (two-sided). Data analyses were done by an investigator who did not acquire cognitive function, MRI or vasoreactivity data.

## Results

### Cognitive function, CBV, CVR and vasoreactivity of mTBI versus sham

There was no difference in NOR or NOL at 10 weeks or 6 months between mTBI and sham rats (Figure 2). At 12 months, NOR but not NOL was lower in mTBI rats versus sham. CBV and CRV were obtained from select brain regions (Figure 1). CBV was higher in the trunk and DG regions of mTBI versus sham rats at 10 weeks but not at 12 months (Figure 3A). There was no difference in CBV in the other regions at both timepoints. CVR was lower in the trunk region in mTBI versus sham rats at 12 months but not at 10 weeks (Figure 3B). The other regions showed no difference between mTBI and sham.

**Figure 2.**
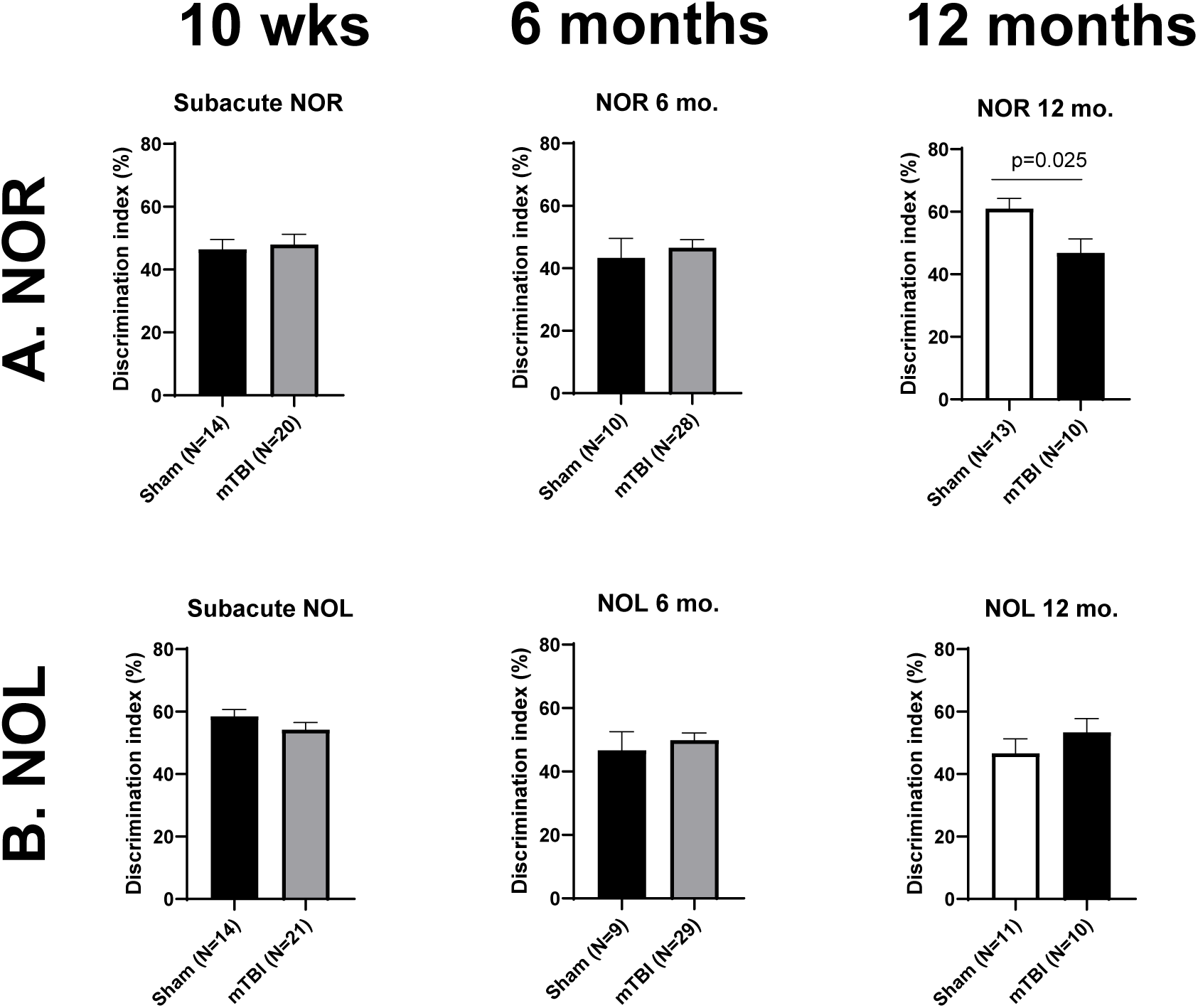
Cognitive function following mTBI. A-B. Following mTBI, there was no difference in novel object recognition (NOR) and novel object location (NOL) at 10 weeks and 6 months post-injury when compared to sham controls. At 12 months, NOR discrimination index was worse in mTBI versus sham, with no difference in NOL.

**Figure 3.**
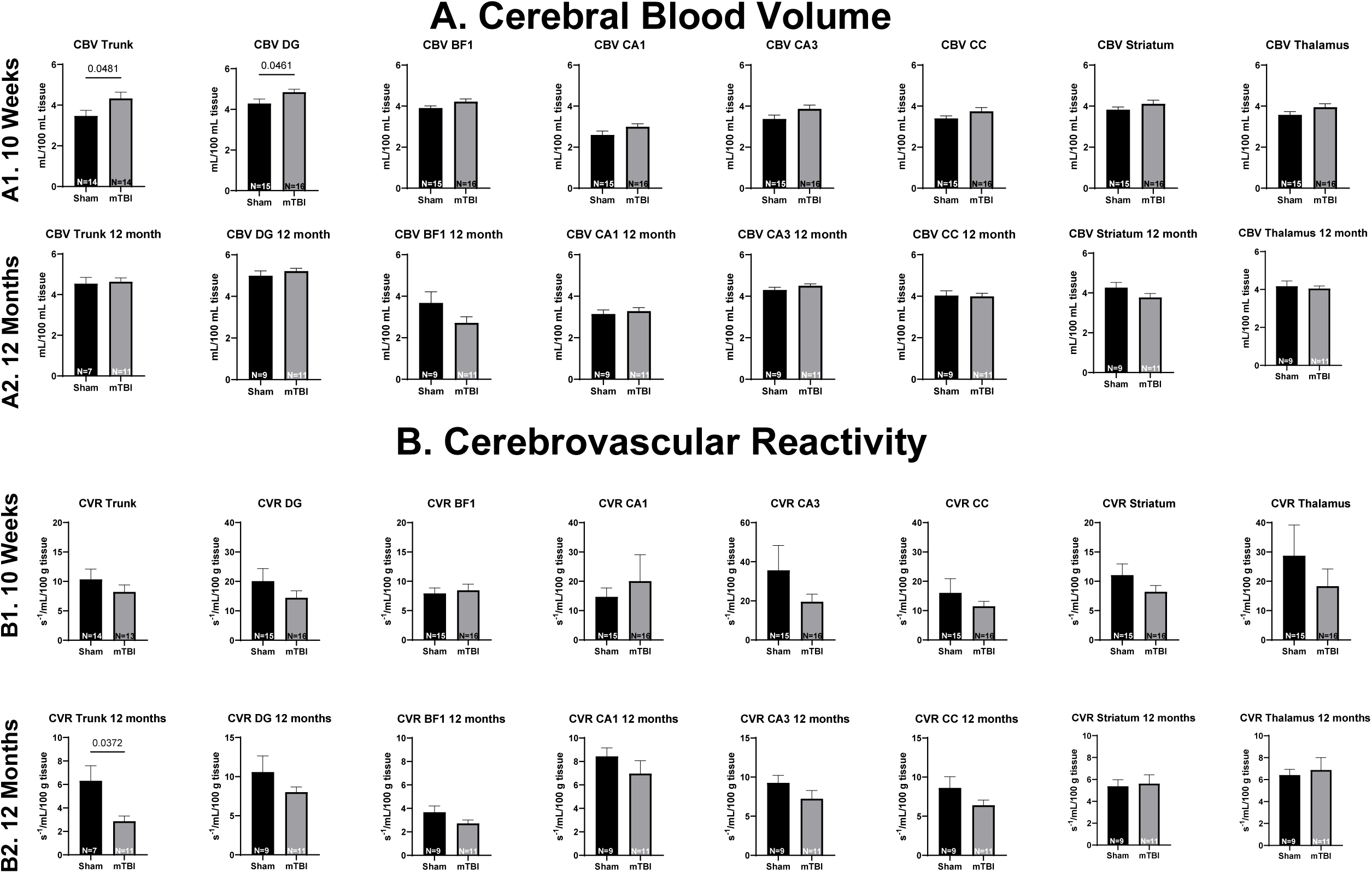
Cerebral blood volume (CBV) and cerebrovascular reactivity (CVR) following mTBI. A1 shows MRI data at 10 weeks demonstrating significantly increased CBV in DG (?) and trunk regions in mTBI versus sham controls. CBV in BF1, CA1, CA3, CC, striatum and thalamus did not differ between the two groups. A2 shows no difference in CBV at 12 months in all regions between sham and mTBI. B1 shows that that cerebrovascular reactivity to hypercarbia exposure at 10 weeks did not differ between mTBI and sham groups. B2 shows that at 12 months, there was lower CVR in trunk region in mTBI versus sham. The rest of the regions did not show any difference.

At 10 weeks, the constriction response of pial arteries to increasing pressure was greater in sham rats (with significant difference at 90 mm Hg intraluminal pressure) versus mTBI, but this difference was not seen at 12 months (Figure 4A-B). Interestingly, the constriction response to intraluminal pressure was significantly higher (90 mm Hg) in pial arteries of sham rats at 10 weeks versus 12 months, whereas the response was similar at 10 weeks and 12 months in mTBI pial arteries (Figure 4C-D). The pial constriction response to angiotensin II and dilator response to NO donor DETA-NONOate showed no difference between mTBI and sham at 10 weeks or 12 months.

**Figure 4.**
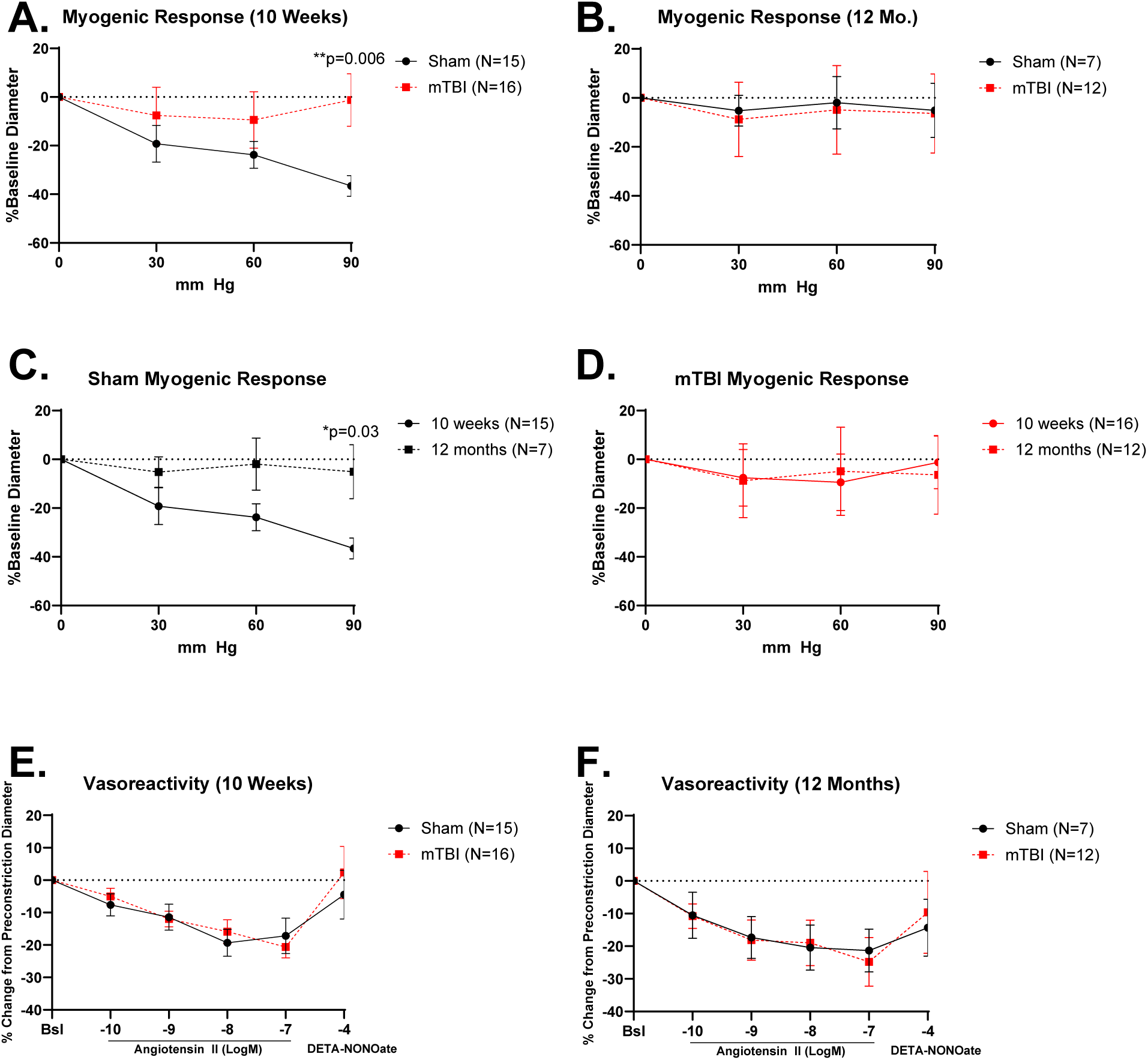
Cerebral arterial vasoreactivity. A-B shows constrictor response to increasing intraluminal pressure at 10 weeks (A) or 12 months (B). There was greater constriction response in sham rats versus mTBI with 90 mm Hg pressure. There was no difference in response at 12 months with blunted constrictor response compared to 10 weeks in sham rats and similar response between 10 weeks and 12 months in mTBI rats. C shows that sham rats have lower pial arterial constrictor response to 90 mm Hg intraluminal pressure at 12 months of study compared to 10 weeks of study, while D shows no difference in mTBI rats at the two timepoints. E-F shows response to Angiotensin II and DETA NONOate. There was progressive cerebral arterial constriction response to increasing doses of angiotensin and dilator response to DETA-NONOate in both sham and mTBI rats at 10 weeks and 12 months with no difference between the two groups.

### Relationship between cognitive function and vascular function outcomes

To assess the relationship between cognitive function and vascular function. Correlation analyses were performed. There was no correlation between regional CBV or regional CVR to NOR or NOL at 10 weeks and 12 months (Table 1). There was also no correlation between pial arterial constriction response to 90 mm Hg intraluminal pressure, angiotensin-II and dilator response to DETA-NONOate and 10 week or 12 month NOR or NOL (Table 2). There was a significant negative correlation between 10 week NOR and pial arterial constriction response to 60 mm Hg intraluminal pressure.

**Table 1.** Correlation between cognitive function score and cerebral blood volume (CBV) and cerebrovascular reactivity (CVR) at 10 weeks and 12 months.

|  | 10 Weeks |  |  |  | 12 Months |  |  |  |
| --- | --- | --- | --- | --- | --- | --- | --- | --- |
|  | NOR |  | NOL |  | NOR |  | NOL |  |
| **CBV** | **R** | **p-value** | **R** | **p-value** | **R** | **p-value** | **R** | **p-value** |
| DG | 0.025 | NS | 0.243 | NS | 0.025 | NS | 0.243 | NS |
| Trunk | 0.257 | NS | 0.238 | NS | 0.257 | NS | 0.238 | NS |
| BF1 | -0.026 | NS | 0.346 | NS | -0.026 | NS | 0.46 | NS |
| CA1 | -0.047 | NS | 0.218 | NS | -0.047 | NS | 0.218 | NS |
| CA3 | 0.125 | NS | 0.031 | NS | 0.125 | NS | 0.031 | NS |
| CC | 0.204 | NS | 0.322 | NS | 0.204 | NS | 0.322 | NS |
| Striatum | 0.194 | NS | 0.344 | NS | 0.194 | NS | 0.344 | NS |
| Thalamus | 0.18 | NS | 0.382 | NS | 0.18 | NS | 0.382 | NS |
| **CVR** |  |  |  |  |  |  |  |  |
| DG | -0.146 | NS | -0.006 | NS | -0.146 | NS | -0.006 | NS |
| Trunk | -0.195 | NS | -0.068 | NS | -0.195 | NS | -0.068 | NS |
| BF1 | -0.215 | NS | -0.152 | NS | -0.215 | NS | -0.152 | NS |
| CA1 | -0.154 | NS | 0.098 | NS | -0.154 | NS | 0.098 | NS |
| CA3 | 0.059 | NS | -0.144 | NS | 0.059 | NS | -0.144 | NS |
| CC | -0.146 | NS | 0.161 | NS | -0.146 | NS | 0.161 | NS |
| Striatum | -0.047 | NS | 0.077 | NS | -0.047 | NS | 0.077 | NS |
| Thalamus | -0.333 | NS | 0.265 | NS | -0.333 | NS | 0.265 | NS |

**Table 2.**
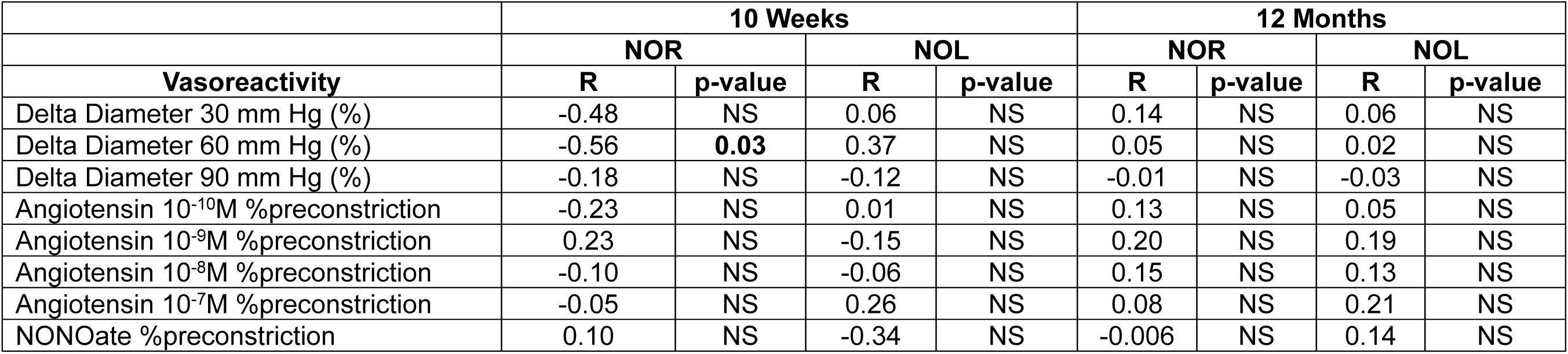
Correlation between cognitive function score and cerebral arterial vasoreactivity at 10 weeks and 12 months.

|  | 10 Weeks |  |  |  | 12 Months |  |  |  |
| --- | --- | --- | --- | --- | --- | --- | --- | --- |
|  | NOR |  | NOL |  | NOR |  | NOL |  |
| **Vasoreactivity** | **R** | **p-value** | **R** | **p-value** | **R** | **p-value** | **R** | **p-value** |
| Delta Diameter 30 mm Hg (%) | -0.48 | NS | 0.06 | NS | 0.14 | NS | 0.06 | NS |
| Delta Diameter 60 mm Hg (%) | -0.56 | **0.03** | 0.37 | NS | 0.05 | NS | 0.02 | NS |
| Delta Diameter 90 mm Hg (%) | -0.18 | NS | -0.12 | NS | -0.01 | NS | -0.03 | NS |
| Angiotensin 10<sup>-10</sup>M %precontraction | -0.23 | NS | 0.01 | NS | 0.13 | NS | 0.05 | NS |
| Angiotensin 10<sup>-9</sup>M %precontraction | 0.23 | NS | -0.15 | NS | 0.20 | NS | 0.19 | NS |
| Angiotensin 10<sup>-8</sup>M %precontraction | -0.10 | NS | -0.06 | NS | 0.15 | NS | 0.13 | NS |
| Angiotensin 10<sup>-7</sup>M %precontraction | -0.05 | NS | 0.26 | NS | 0.08 | NS | 0.21 | NS |
| NONOate %precontraction | 0.10 | NS | -0.34 | NS | -0.006 | NS | 0.14 | NS |

### Effects of exercise on cognitive and vascular function post-mTBI

A separate group of mTBI rats underwent early (2 weeks post-injury) or late (10 months post-injury) exercise to evaluate effects of exercise on cognitive and vascular function post-mTBI. Over the span of 6 weeks of exercise, the total distance traveled was 5896±767 m (early) and 6968±826 m (late) (p=NS). Although there were overall group differences in NOR at 12 months (ANOVA p=0.03), post-hoc pairwise analysis did not show a statistically significant difference between each group (Figure 5A). There was a trend (not significant) of increase in NOR in mTBI with early exercise versus mTBI, but not with late exercise. There was no difference in NOL among the groups (Figure 5B).

**Figure 5.**
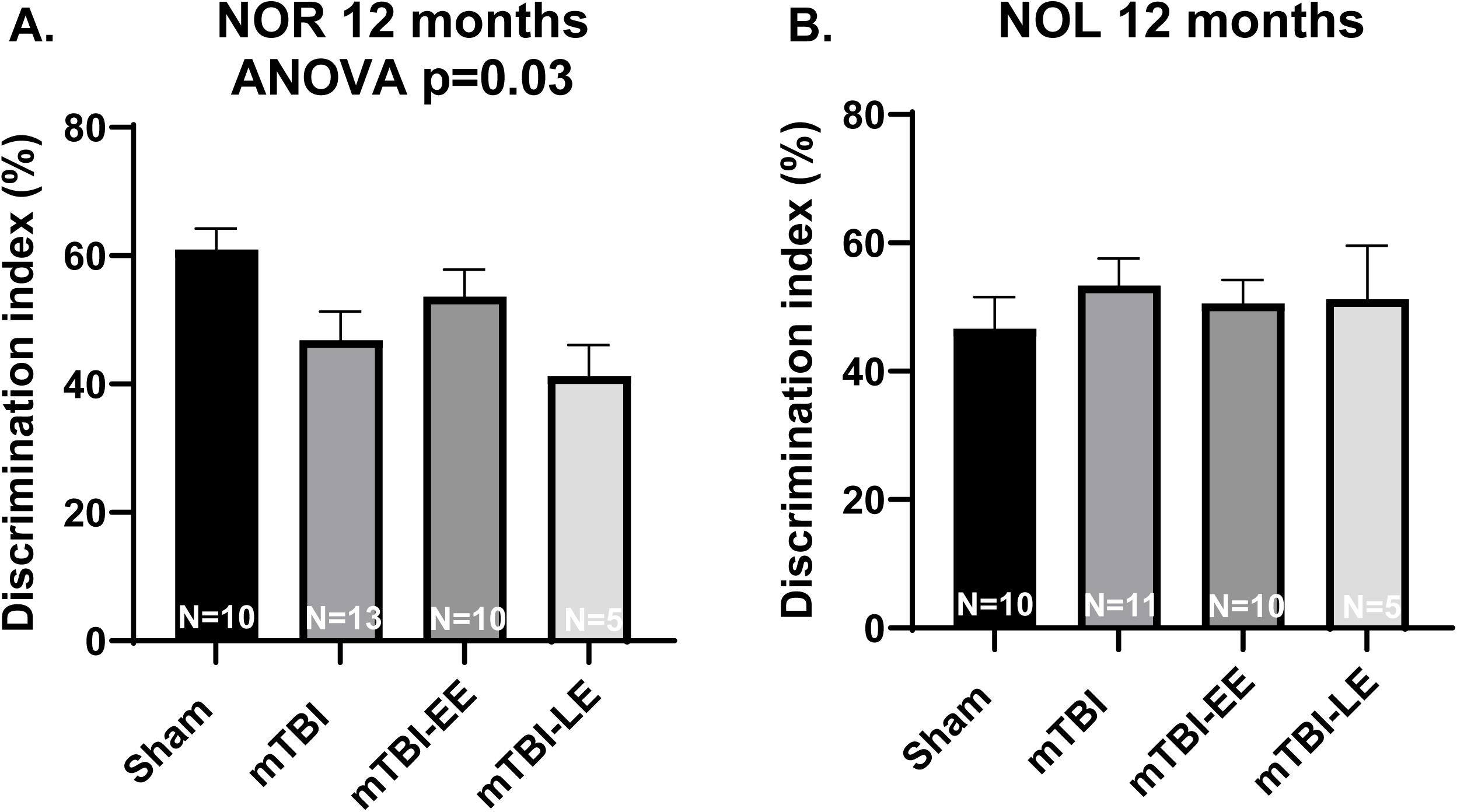
Cognitive function following mTBI without or with aerobic exercise. A shows overall significant group differences in NOR at 12 months among sham, mTBI, mTBI with early exercise and mTBI with late exercise. Despite trend towards increased NOT in mTBI with early exercise versus mTBI, the pairwise post-hoc analyses showed no significant differences in NOR between mTBI without and with early or late exercise. B shows no difference in NOL among the groups.

There were no significant differences in pial arterial constriction response to intraluminal pressure, angiotensin II or dilator response to DETA-NONOate among sham, mTBI, mTBI with early exercise or mTBI with late exercise (Figure 6 A-B). Similarly, no difference was observed in the same groups as regards regional CBV and regional CVR at 12 months (Figure 7 A-B).

**Figure 6.**
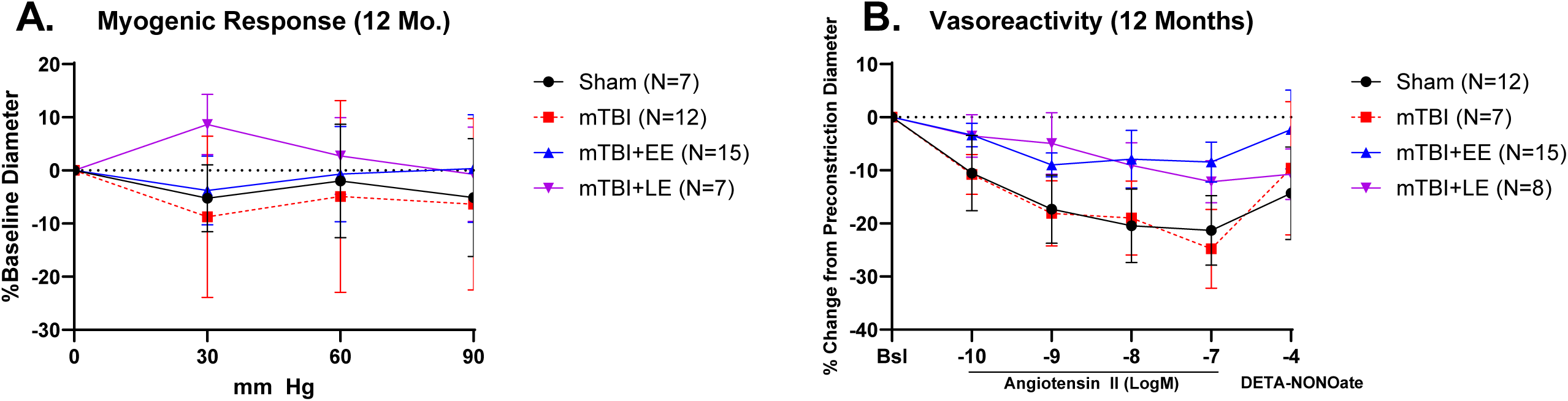
Cerebral arterial vasoreactivity following mTBI without or with aerobic exercise. A shows no difference in 12 month pial arterial constrictor response to increasing intraluminal pressure among sham, mTBI, mTBI with early exercise and mTBI with late exercise. B shows similar results with constrictor response to angiotensin II and dilator response to DETA-NONOate.

**Figure 7.**
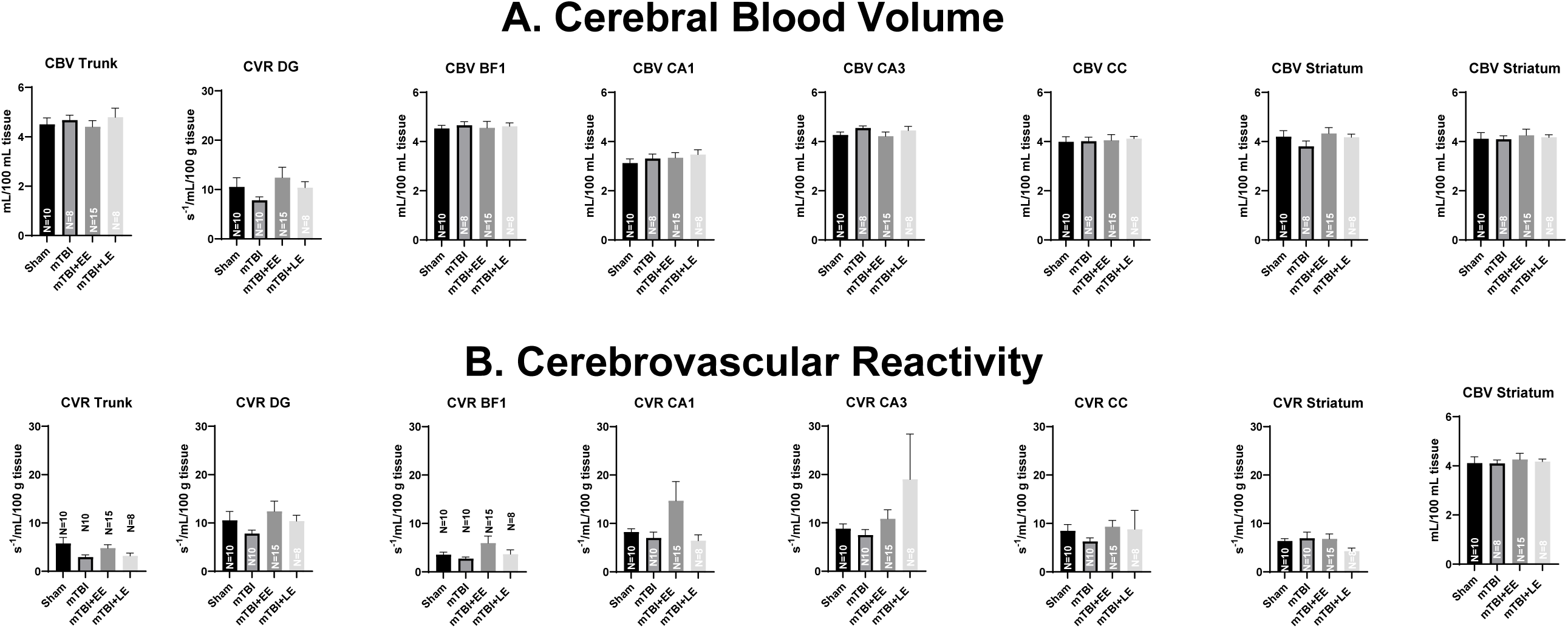
Cerebral blood volume (CBV) and cerebrovascular reactivity (CVR) following mTBI without or with aerobic exercise. Regional CBV (A) and CVR (B) were not significantly different among sham, mTBI rats without or with early or late exercise.

## Discussion

We present the first comprehensive simultaneous assessment of cognitive and cerebrovascular dysfunction at 12 months following mTBI in Sprague-Dawley rats that we are aware of and report the effects of aerobic exercise in modulating neurovascular outcomes. The findings demonstrate chronic persistent cognitive dysfunction at 12 months mainly in the short-term memory domain, impaired cerebrovascular myogenic response at 10 weeks and regional impaired vasoreactivity to hypercapnea at 12 months following mTBI in rats. The association between short-term memory impairment and cerebrovascular impairment was not demonstrated. Although a trend towards improved short-term memory in mTBI rats that underwent early exercise versus mTBI rats without aerobic exercise was seen, the difference was not significant and there is no definitive evidence that early or late exercise improves cognitive or vascular function following mTBI.

Preclinical studies show that exposure to severe TBI lead to late (1-year) development of cognitive impairment^7,36,37^ but there is a dearth of evidence in preclinical models that mTBI also develop similar pathology. This is a critical gap since mTBI represents the great majority of TBI cases with estimates ranging from 60-95% of all cases^20,21^. There is no established consensus classification scheme for TBI severity, and the definition of mTBI, which includes the common concussion, has been a matter of controversy^20,38^. Despite this, the FPI model used in this study was found to be consistent with the mTBI classification schemes used by either the United States Department of Defense or Veterans Affairs classification^22,39^. The FPI model leads to non-catastrophic TBI with acute physiological disruption followed by recovery within a few days and absence of gross histopathological damage and lack of cavitation months post-injury^22,40,41^. The current study shows that mTBI leads to impairment at 1 year in the cognitive domain associated with short-term memory (tested by NOR) but not long-term spatial memory (tested by NOL). Our prior study showed impaired NOR at 6 months following mTBI^22^ but surprisingly, our current cohort showed impairment at 12 but not at 6 months. NOL and NOR are simple behavioral assays of memory that rely on the rat’s innate exploratory behaviors and preference for novelty^42^. NOL primarily evaluates spatial learning which relies heavily on hippocampal activity while the NOR evaluates non-spatial learning which relies on multiple brain regions. NOR is a commonly used task to evaluate general memory function in rodents with some studies suggesting that hippocampal lesions or inactivation do not affect NOR while others show the opposite^43^. The NOR abnormality we observed at 12 months may be consistent with the diffuse injury induced by the FPI model which affects multiple brain regions. Mechanical forces of the fluid pulse reflect off the skull temporal ridge and affect diffuse regions such as the hippocampal area CA3, primary somatosensory barrel cortex and ventral posterior nuclei of the thalamus^22^. In a large study comparing patients with mTBI versus no TBI exposure, mTBI was associated with increased risk of developing dementia with a hazard ratio of 3.8 (95% confidence interval 2.1-6.9)^14^. AD dementia was shown to have significant impairment in memory tests involving both novel object recognition and object locations^44^ and our 12 month preclinical observation of novel object recognition-associated cognitive dysfunction is likely modeling observations in mTBI patients of greater predisposition of later development of dementia.

Cerebrovascular dysfunction is a well-known effect of severe, acute TBI with ischemic brain damage evident at autopsy in >90% of acute TBI mortalities^22,45^. In contrast, the chronic cerebrovascular effect of mTBI is less well understood. mTBI patients were found to have 2.2 times increased risk of ischemic stroke and transient ischemic attacks compared to matched controls without TBI^14^ through still undefined mechanisms. Preclinical data show that TBI acutely led to reduced local cerebral blood flow and concomitant neurovascular uncoupling^46^. Our prior work showed no difference in regional CBV and cerebral blood flow by MRI between sham and mTBI rats at 6 months but there was altered angiotensin II-stimulated smooth muscle-dependent vasoreactivity in mTBI rats^22^. In the current work, we again utilized *in vivo* MRI readouts of cerebrovascular function by measuring regional CBV but in addition, we now added CRV, which measures change in regional blood flow following hypercapneic insult, a measure of dynamic cerebrovascular autoregulation. We also added *ex-vivo* investigation of circle of Willis (pial artery) response to increased intraluminal pressure and tested vascular smooth muscle contractile response to increasing doses of angiotensin II and dilator response to NO donor (DETA-NONOate). Our *in vivo* MRI data showed a consistent pattern of increased CBV in various brain regions in mTBI versus sham rats at 10 weeks although only the trunk primary somatosensory cortex and dentate gyrus regions showed significant differences. No difference in CBV was noted between mTBI and sham at 12 months. On the other hand, CVR was lower in the trunk region of mTBI versus sham at 12 months. We found no significant correlation between cognitive function test outcomes (NOR and NOL) and MRI vascular outcomes (CBV and CVR). Notably, as previously described, we found impaired NOR in mTBI versus sham rats at 12 months. In human studies, blood oxygenation level-dependent response using functional MRI showed greater activation in the temporooccipital and frontal cortical areas specifically during object recognition behavior^47,48^. These regions include the trunk primary somatosensory cortex, and the impaired CVR in this region could potentially be linked to the short-term memory impairment observed in mTBI rats.

We did not find any difference in cerebrovascular response to angiotensin II or DETA-NONOate between mTBI and sham rats either at 10 weeks or at 12 months. We did have an interesting finding of significant reduction in constriction response to increased intraluminal pressure (myogenic response) in mTBI cerebral arteries versus sham at 10 weeks but the blunted response to intraluminal pressure was observed in both sham and mTBI rats at 12 weeks. The temporal change in response in sham rats suggests that the aging cerebral artery loses myogenic response to intraluminal pressure, while the response seen with mTBI suggests that this impaired myogenic response is acquired 10 weeks following injury and persists to 12 months. This observation could suggest that mTBI confers an early aging phenotype in terms of cerebrovascular myogenic response. Whether this early impairment is modulating the observed cognitive dysfunction and impaired CVR in the trunk region at 12 months remains unsettled by this study but should be the focus of future investigation.

There is no established treatment to mitigate the long-term adverse cognitive and vascular function effects of TBI. Vigorous aerobic exercise training in patients with chronic TBI resulted in improved cognitive function scores strongly related to gains in cardiovascular fitness^49^. Aerobic exercise training was shown to improve the autoregulatory domains controlling cerebrovascular function. Increased carbon dioxide production from aerobic exercise leads to greater cerebral vasoreactivity to regulate flow in hypercapnea or hypocapnia^50^. Increases in systemic pressure with low intensity exercise is counterregulated by cerebral autoregulation^51^. Additionally, sustained muscle engagement during exercise leads to cortical activation in motor and sensorimotor areas increasing cerebral metabolism and engaging neurovascular coupling^52^. In the rehabilitation of TBI patients, aerobic exercise was shown to be safe after the acute recovery period^53^. We tested two approaches to aerobic exercise following mTBI, an early exercise regimen starting at 2 weeks following injury (immediately following acute recovery) and late exercise regimen (10 months following injury). Results did not demonstrate a statistically significant change with either early or late exercise in terms of cognitive function and vascular function outcomes, although there was a trend towards improved 12 month NOR and trunk CVR with early exercise. Future studies should further explore this relationship utilizing greater sample size with greater power to tease a difference.

The study has several important limitations. We limited our cognitive function assays to NOR and NOL which tests two cognitive domains. The choice of these tests is based on balancing the assessment of multiple relevant cognitive-behavioral domains while ensuring practical logistical feasibility to successfully complete the assays within a tight time window. It is possible that other cognitive domains are also impaired and that these may be affected by cerebrovascular dysfunction. In the future, other tests such as the Morris water maze, Barnes maze or radial arm maze could be utilized. The MRI vascular outcome readouts were also done at one timepoint prior to terminal sacrifice missing out on information that could be provided by longitudinal follow up of vascular changes. We initially planned a graduated weekly increase in aerobic exercise speed, but some rats could not tolerate the increased speed so we kept the maximum speed that each rat could tolerate to maintain aerobic exercise. A more prolonged exercise regimen given this limitation may be explored in future studies to assess for benefit. Sample size limitations may have limited the power of the study to tease out differences, and our current data could be used to inform sample size computations of future studies.

In conclusion, 12 months following mTBI, rats developed impaired short term memory cognitive function and cerebrovascular reactivity to hypercapnea in the trunk primary somatosensory cortex region compared to sham controls. Sham rats demonstrated impaired myogenic response at 12 months when compared to 10 weeks, while the onset of this impairment in mTBI rats was early at 10 weeks. We found no correlation between cognitive and vascular function outcomes. Early or late exercise did not significantly affect the cognitive and vascular impairments observed at 12 months. These physiologic findings enhance our understanding of the mechanisms underlying the long-term neurovascular pathologies following mTBI.

## Acknowledgments

We thank Miss Gail Farrell for assistance with regulatory compliance and the Phoenix VA Office of Research and Arizona Veterans Research and Education Foundation for management and logistical support. The content of this manuscript does not represent the views of the US Department of Veterans Affairs or the United States government.

